# Arabidopsis ROOT PHOTOTROPISM2 is a Light-Dependent Dynamic Modulator of Phototropin1

**DOI:** 10.1101/862649

**Authors:** Taro Kimura, Tomoko Tsuchida-Mayama, Hirotatsu Imai, Koji Okajima, Kosuke Ito, Tatsuya Sakai

## Abstract

*Arabidopsis thaliana* phototropin1 (phot1) is a blue-light photoreceptor, i.e. a blue-light-activated Ser/Thr-protein kinase that mediates various light responses including phototropism. Phot1 functions in hypocotyl phototropism dependent on the light induction of ROOT PHOTOTROPISM2 (RPT2) proteins within a broad range of blue light intensities. It is not yet known however how RPT2 contributes to the photosensory adaptation of phot1 to high intensity blue light and the second positive phototropism. We here show that RPT2 suppresses the activity of phot1. Yeast two-hybrid analysis indicated RPT2 binding to the LOV1 (light, oxygen or voltage sensing 1) domain of phot1 required for its high photosensitivity. Our biochemical analyses revealed that RPT2 inhibits the autophosphorylation of phot1, suggesting that it suppresses the photosensitivity and/or kinase activity of phot1 through the inhibition of LOV1 function. We found for the first time that RPT2 proteins are degraded via a ubiquitin-proteasome pathway when phot1 is inactive and stabilized under blue-light conditions in a phot1-dependent manner. We propose that RPT2 is a molecular rheostat that maintains a moderate activation level of phot1 under any light intensity conditions.

## INTRODUCTION

Plant life is strongly dependent on light and the angiosperm *Arabidopsis thaliana* uses several kinds of photoreceptors to effectively adapt to various light conditions, including phytochromes (phys) which are red-/far-red-light photoreceptors, cryptochromes (crys), phototropins (phots), the LOV (light, oxygen or voltage sensing)/F-box/Kelch-domain proteins (ZTL, FKF1, and LKP2), which are blue-light photoreceptors, and a UV photoreceptor UVR8 (de Wit et al., 2016). In the case of the phy and cry families, phyA and cry2 are highly expressed in *Arabidopsis* seedlings under dark conditions and become unstable under bright light conditions, thus contributing strongly to the responses to weak and not bright light conditions. (Clough and Vierstra, 1997; Lin et al., 1998; Sharrock and Clack, 2002; Casal et al., 2014). On the other hand, phyB and cry1 are stable and function as major photoreceptors under bright light conditions (Lin et al., 1998; Li et al., 2011). Plants therefore use different photoreceptor families and members of these families to recognize light quality and quantity in order to adapt to their various light environments.

In the case of the phot family, the highly photosensitive photoreceptor phot1 and the less photosensitive photoreceptor phot2 function redundantly in a fluence-rate dependent manner (Sakai et al., 2001). The phots have LOV1 and LOV2 domains in their N-terminal portions, and a serine/threonine (Ser/Thr) kinase domain belonging to the AGC (for cAMP-dependent protein kinase A, cGMP-dependent protein kinase G, and phospholipid-dependent protein kinase C) VIII kinase domain, within their C-terminal half (Christie, 2007; Rademacher and Offringa., 2012). The phots localize on the inner surface of the plasma membrane, show autophosphorylation activities under blue-light conditions, and mediate blue-light responses such as phototropism, chloroplast photorelocation and stomatal opening (Christie et al., 1998; Kagawa et al., 2001; Kinoshita et al., 2001; Sakai et al., 2001; Sakamoto and Briggs, 2002). Each LOV domain harbors flavin mononucleotide (FMN) as a blue-light absorbing chromophore (Christie et al., 1998; Sakai et al., 2001) and transiently forms a cysteinyl adduct with a blue-light-excited FMN (Christie, 2007). This cysteinyl adduct undergoes thermal decay and LOV domains thus show dark reversion (Okajima, 2016). The photochemical reaction mediated through the LOV2 domain is indispensable for phot function and leads to their conformational change and activation as a Ser/Thr kinase (Christie et al., 2002; Christie, 2007). The lifetime of the cysteinyl-FMN adduct of phot2 LOV2 is 10-fold greater than that of phot1, which appears to be one of reasons why phot2 does not function under low intensity blue light conditions (Okajima et al., 2012). On the other hand, phot1 is a unique photoreceptor that mediates the phototropic responses in etiolated hypocotyls over a broad dynamic range of blue light intensities between 10^−5^ and 10^2^ µmol m^−2^ s^−1^ (Sakai et al., 2001; Haga et al., 2015). The photochemical reaction at the LOV1 of phot1 is unnecessary for the phototropic response (Christie et al., 2002), but this domain itself is necessary for the induction of the phototropic responses under low intensity blue light conditions (Sullivan et al., 2008). The phot1 LOV1 domain thus appears to play an important role in the control of phot1 photosensitivity, but the precise underlying molecular functions are yet unrevealed.

The ROOT PHOTOTROPISM2 (RPT2) protein is a signal transducer in *Arabidopsis* phototropism (Sakai et al., 2000). It localizes on the plasma membrane and forms a complex with phot1 in vivo (Inada et al., 2004). RPT2 expression is suppressed in etiolated seedlings and upregulated by red- and/or blue-light irradiation (Sakai et al., 2000; Tsuchida-Mayama et al., 2010). The phys and crys are necessary for the induction of *RPT2* transcription (Tsuchida-Mayama et al., 2010). *Rpt2* loss-of-function mutants exhibit increased responses to very low intensity blue light at 10^−5^ µmol m^−2^ s^−1^ and decreased responses to blue light at 10^−3^ µmol m^−2^ s^−1^ or more during hypocotyl phototropism (Haga et al., 2015). On the other hand, the expression of *RPT2* prior to a phototropic stimulation in etiolated wild-type seedlings accelerates continuous light-induced phototropism (Haga et al., 2015). These findings have suggested that the light induction of RPT2 expression reduces the photosensitivity of phot1, which is required for the photosensory adaptation of phot1 and the second phototropism under bright light conditions (Haga et al., 2015).

In our present study, we tested the hypothesis that RPT2 controls the photosensitivity of phot1 through its LOV1 domain. A yeast two-hybrid assay indicated that RPT2 binds to the phot1 LOV1 domain and immunoblotting using Phos-tag SDS-poly-acrylamide gels indicated that a *rpt2* mutation enhances the autophosphorylation of phot1, and that the *RPT2* overexpression suppresses this. These data indicated that RPT2 controls the autophosphorylation activity of phot1 through the LOV1 domain. We further showed that RPT2 expression is upregulated not only by the phys and crys but also by phots. Based on our current results, we propose that RPT2 acts as a molecular rheostat that maintains a moderate activation of phot1 under any light intensity conditions.

## RESULTS

### RPT2 binds to the LOV1 domains of phot1

Our previous study demonstrated that the N-terminal half of RPT2 (RPT2 N) including the BTB/POZ (broad complex, tramtrack and bric-à-brac/Pox virus and zinc finger) protein-protein interaction domain binds to the N-terminal half of PHOT1 that includes two LOV domains (Inada et al., 2004). We divided the N-terminal half of PHOT1 into 4 fragments and examined RPT2 N binding to these fragments or its kinase domain (PHOT1 C) using a yeast two-hybrid assay (Figure 1A: Inada et al., 2004). As shown in Figure 1B, RPT2 N bound only to the LOV1 domain (PHOT1 N2). We confirmed its binding using an in vitro pull-down assay. The hemagglutinin-tagged (HA-) RPT2 N proteins specifically interacted with the histidine/ProS2-tagged (His-) PHOT1 N2 proteins on metal affinity resins in contrast with His-PHOT1 N4 harboring the LOV2 domain (Figure 1C). These results suggest that RPT2 affects phot1 photosensitivity via the LOV1 domain.

**Figure 1.**
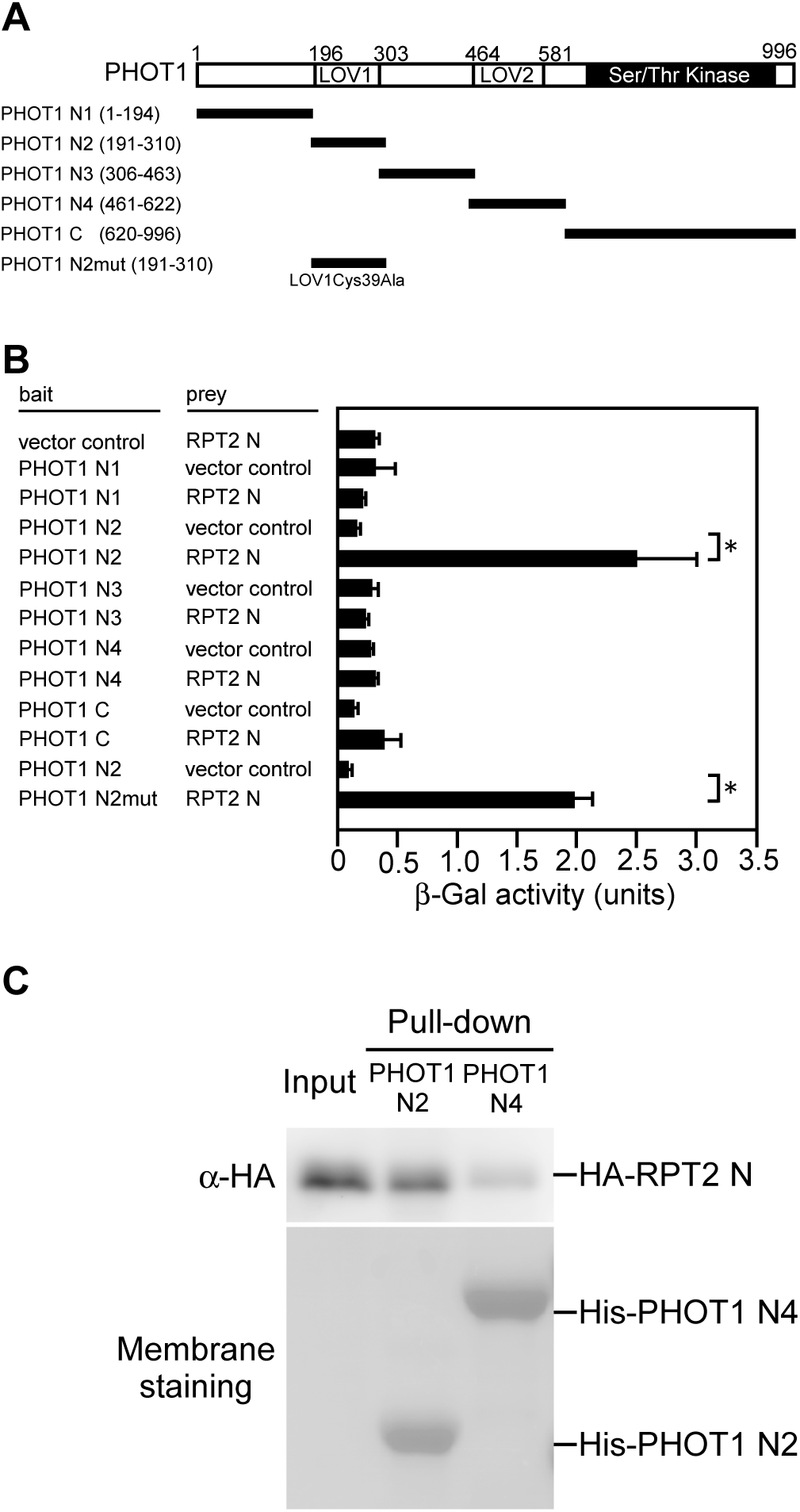
Interaction between phot1 and RPT2. **(A)** Schematic representation of the phot1 structure. The Ser/The protein kinase and LOV domains are denoted by the solid and dotted blocks, respectively. The amino acid residues used for each underlined construct are indicated in parentheses. **(B)** Yeast two-hybrid assay of phot1-RPT2 interactions. Solution assays of β-Gal activity were performed for the combinations indicated on the left. The data shown are means values ± SE (*n* = 3). The asterisk denotes statistically significant differences compared with the vector control (Student’s t test; *P* < 0.05). **(C)** In vitro pull-down assay to verify the interaction between the LOV domains of phot1 and the N-terminal half of RPT2. HA-tagged proteins of N-terminal half of the RPT2 (HA-RPT2 N) were incubated with the His-tagged LOV1 or LOV2 domains of phot1 for His-tag pull-down assay and detected by immunoblotting using anti-HA (α-HA) antibodies. The protein-blotted membrane were stained using a Pierce reversible protein staining kit.

Christie et al. (2002) reported previously that by using the LOV1Cys39Ala mutant, the LOV1 domain of phot1 is photochemically active but that its photochemical reaction is unessential for phot1 function. Our current yeast two hybrid assay data showed that the LOV1 domain harboring a Cys39Ala mutation (PHOT1 N2mut) also binds to RPT2 N as well as PHOT1 N2 (Figure 1B). Furthermore, the *phot1 phot2* transgenic seedlings expressing the *PHOT1* gene with the LOV1Cys39Ala mutation showed a shortened time lag to the induction of phototropic responses by a red-light pretreatment (Supplemental Figure 1), which is dependent on *RPT2* (Haga et al., 2015). Our previous study had already indicated that RPT2 can form a complex with phot1 in vivo under both conditions of darkness and blue light (Inada et al., 2004). These results suggest that RPT2 binding and function are unaffected by the photochemical reaction of the phot1 LOV1 domain and the phosphorylation state of phot1.

### RPT2 suppresses the autophosphorylation of phot1

Our previous genetic study suggested that RPT2 reduces the photosensitivity of phot1, which is required for a second positive phototropism under bright light conditions (Haga et al., 2015). This indicated that RPT2 may suppress the activity of phot1. To test this possibility, we examined the autophosphorylation pattern of phot1 in both wild-type *Arabidopsis* seedlings and *rpt2* mutants grown on the surface of vertically oriented agar medium. We conducted immunoblotting analysis using a Phos-tag acrylamide gel for this experiment (Kinoshita and Kinoshita-Kikuta, 2011), in which the migration of phosphorylated proteins is specifically retarded. We observed autophosphorylation patterns of phot1 in response to blue-light irradiation for 2 h at 0.001, 0.1 and 100 µmol m^−2^ s^−1^ (Figure 2A). When the wild-type seedlings were irradiated, the mobilities of PHOT1 proteins became much slower if the fluence rates were higher (Figure 2A). These mobility shifts were more clearly observed using a Phos-tag acrylamide gel (+Phos-tag) compared with a normal SDS-polyacrylamide gel (– Phos-tag: Figure 2A). This result suggested that the autophosphorylation activity of phot1 increases as the fluence rates increase.

**Figure 2.**
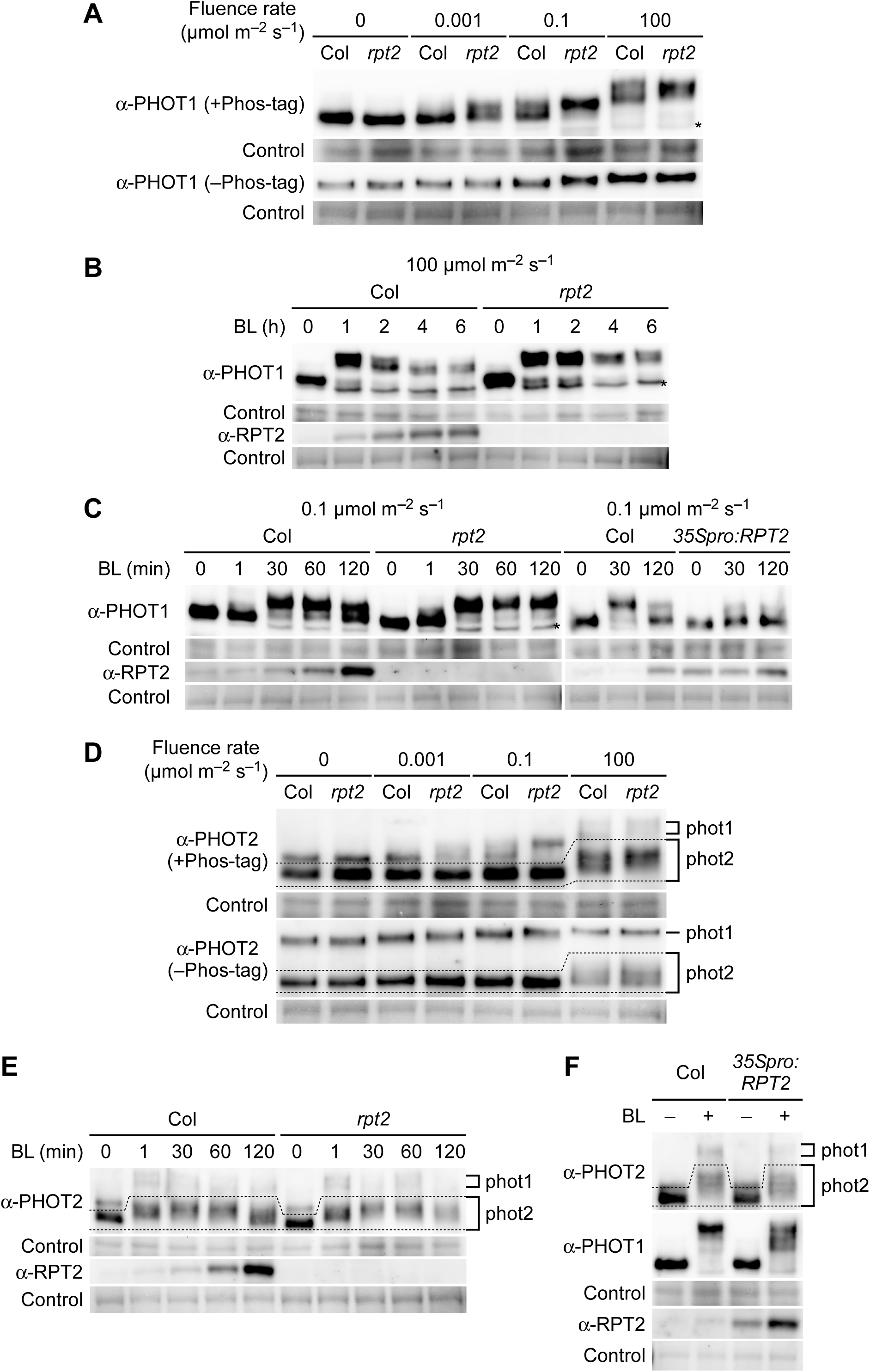
RPT2 suppresses the autophosphorylation of phot1. **(A)** Immunoblotting analysis of PHOT1 in wild type (Col) and *rpt2* mutant *Arabidopsis*. Two-day-old etiolated seedlings were irradiated with unilateral blue light at the indicated fluence rates for 2 h. Total proteins (20 µg) extracted from the seedlings were separated on 6% SDS-PAGE gels with 2 µM Phos-tag (+ Phos-tag), followed by immunoblotting with anti-PHOT1 (α-PHOT1) antibodies. 10 µg of the total proteins were separated on 6% SDS-PAGE gels without Phos-tag (– Phos-tag) for comparison. An asterisk indicates a non-specific band. The protein-blotted membranes were stained as a loading control. **(B and C)** Time course analysis of phot1 autophosphorylation. Two-day-old etiolated seedlings were irradiated with unilateral blue light at 100 µmol m^−2^ s^−1^ in **(B)** or 0.1 µmol m^−2^ s^−1^ in **(C)** for the indicated times. Other details are as described in **(A)**. **(D)** Immunoblotting analysis of PHOT2 in wild type (Col) and *rpt2* mutant. Immunoblotting was performed with an anti-PHOT2 antibody (α-PHOT2) antibodies. Other details were as in **(A)**. **(E)** Time course analysis of phot2 autophosphorylation. Two-day-old etiolated seedlings were irradiated with unilateral blue light at 100 µmol m^−2^ s^−1^ for the indicated times. Immunoblotting was performed with α-PHOT2 and anti-RPT2 (α-RPT2) antibodies. Other details are as described in **(A)**. **(F)** Autophosphorylation of phot1 and phot2 in the *rpt2* mutant transformed with a *35Spro:RPT2* gene (*35Spro:RPT2*). Two-day-old etiolated seedlings were irradiated with unilateral blue light at 100 µmol m^−2^ s^−1^ for 30 minutes. Immunoblotting was performed with α-PHOT1, α-PHOT2 and α-RPT2 antibodies. Other details are as described in **(A)**.

We next investigated the autophosphorylation pattern of phot1 during blue light irradiation. When we monitored the phosphorylation status of the PHOT1 protein under blue light conditions at 100 µmol m^−2^ s^−1^ (Figure 2B), mobility shifts of this protein were detectable at 1 hour, but further irradiation suppressed phot1 autophosphorylation in parallel with the accumulation of RPT2 proteins. Under blue light conditions at 0.1 µmol m^−2^ s^−1^, mobility shifts in the PHOT1 proteins were marginally detectable at 1 min, became saturated at 30 min, but were suppressed at 60 and 120 min after the onset of irradiation in wild-type etiolated seedlings (Figure 2C).

*Arabidopsis* wild-type hypocotyls show a delayed phototropic response when grown along the surface of vertically oriented agar medium and a quick response when grown on the agar medium without touching the agar medium (Haga and Sakai, 2012; Sullivan et al., 2019). We examined the autophosphorylation of phot1 in wild-type *Arabidopsis* seedlings grown on agar medium in 0.2 ml tubes (Haga et al., 2012). They showed similar phot1 autophosphorylation patterns to those in the seedlings grown on vertically oriented agar medium (Supplemental Figure 2), suggesting that the phot1 autophosphorylation activity in seedlings is unaffected by the friction or the moisture between the agar surface and the shoots. We therefore used the seedlings grown on the surface of vertically oriented agar medium in later analyses.

In the –Phos-tag gel, the *rpt2* mutation did not produce any obvious effect on the mobility shifts of the PHOT1 proteins (Figure 2A), as reported previously (Inada et al., 2004; Haga et al., 2015). However, in the +Phos-tag gel, the *rpt2* mutation appeared to cause a pronounced mobility shift in the PHOT1 protein under blue light conditions at 0.001, 0.1 and 100 µmol m^−2^ s^−1^ (Figure 2A). The mobility shift of PHOT1 with the *rpt2* mutation disappeared when the extracted proteins were treated with alkaline phosphatase (Supplemental Figure 3), indicating that its shift reflects differences in phosphorylation state of PHOT1. Interestingly, the *rprt2* mutants did not exhibit any attenuation of phot1 autophosphorylation at 2, 4 and 6 hours (Figure 2B) or at 60 and 120 min (Figure 2C) after the onset of blue-light irradiation of 100 and 0.1 µmol m^−2^ s^−1^, respectively. On the other hand, a constitutive *RPT2* expression line (*35Spro:RPT2*: Haga et al., 2015) showed an attenuation of phot1-autophosphorylation at 30 min after the onset of the blue-light irradiation (Figure 2C). These results suggested that RPT2 proteins can suppress the autophosphorylation or enhance the dephosphorylation of phot1 and that a loss-of-function mutation in *RPT2* leads to a continuous hyperactivation of phot1 in seedlings.

The effect of RPT2 on the autophosphorylation of phot2 was also examined using an anti-PHOT2 antibody, which recognizes both PHOT1 (∼120 kDa) and PHOT2 (∼110 kDa: Supplemental Figure 4). When wild-type seedlings were irradiated with blue light at 0.001, 0.1, and 100 µmol m^−2^ s^−1^, mobility shifts in the PHOT2 proteins were detectable only at 100 µmol m^−2^ s^−1^ (Figure 2D, dotted area). Under blue-light conditions of 100 µmol m^−2^ s^−1^, the mobility shifts of the PHOT2 proteins were marginally detectable at 1 min, saturated at 30 min, and attenuated at 120 min after the onset of the irradiation in wild-type etiolated seedlings (Figure 2E). Those autophosphorylation patterns were also detected in the *rpt2* mutants and the *35Spro:RPT2* transgenic lines (Figure 2E and 2F). Although green fluorescent protein (GFP)-fused phot2 formed a complex with RPT2 in vivo (Supplemental Figure 5A and 5B) and RPT2 N showed binding activity to the phot2 LOV1 domain in yeast (Supplemental Figure 5C), the results of our phenotypic analysis suggested that RPT2 has no significant impact on the autophosphorylation of phot2. This finding was consistent with the finding of a previous study that the *rpt2* mutation has no effect on phot2-dependent phototropic responses (Inada et al., 2004).

We next examined the inhibitory effects of RPT2 on the autophosphorylation activity of phot1 using an in vitro phosphorylation assay (Figure 3). This assay was performed with microsomal proteins extracted from *rpt2* mutants transformed with a *pMDC7-RPT2* construct, in which the expression of RPT2 is inducible by estradiol (Est) treatment (Supplemental Figure 6: Zuo et al., 2000). The autophosphorylation of phot1 was detected as a radiolabeled 120 kDa protein in the microsomal fraction as described previously (Liscum and Briggs, 1995). As expected, blue-light irradiation caused the phosphorylation of these 120 kDa proteins in the microsomal fractions of the *pMDC7-RPT2* seedlings and *pER8* vector control line (Figure 3A). On the other hand, Est treatments suppressed this phot1 phosphorylation in the *pMDC7-RPT2* seedlings but not in the *pER8* vector control line (Figure 3). Our immunoblotting analysis confirmed that PHOT1 protein expression was comparable among all of the microsomal fractions and that the RPT2 proteins were expressed only in the microsomal fraction of Est-treated *pMDC7-RPT2* seedlings (Supplemental Figure 6B). These results suggested that Est-induced RPT2 proteins suppress the in vitro autophosphorylation of phot1.

**Figure 3.**
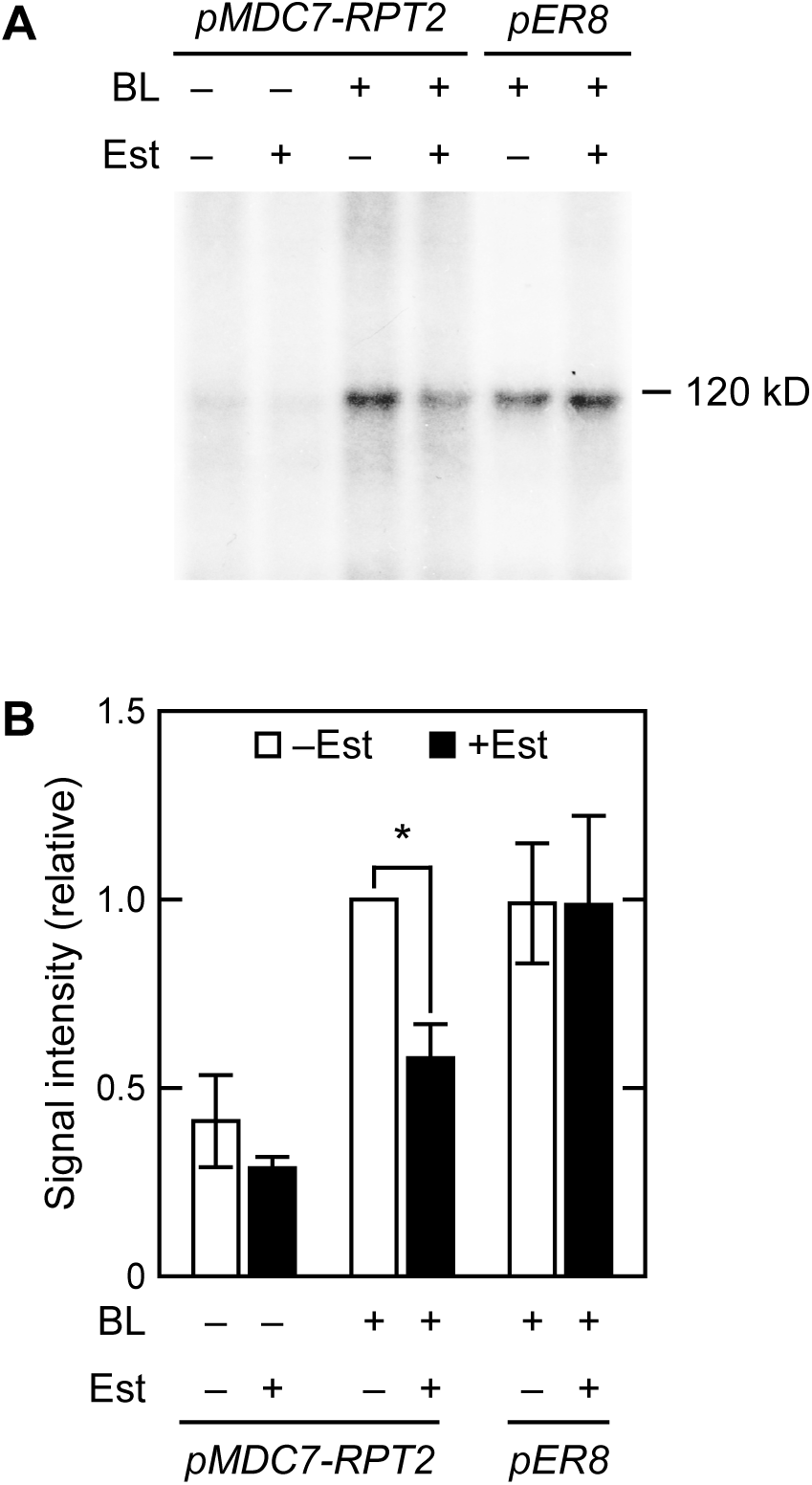
In vitro blue light-induced phosphorylation of a 120-kDa protein in microsomal membranes. Two-day-old etiolated *rpt2* mutants transformed with a *pMDC7-RPT2* construct (*pMDC7-RPT2*) or wild-type seedlings transformed with a *pER8* vector control (*pER8*) were grown on agar medium with (–) or without (+) 10 µM estradiol (Est) under darkness. Microsomal proteins were then extracted. **(A)** In vitro blue light-induced phosphorylation of a 120-kDa protein in microsomal membranes. Microsomal proteins were irradiated with mock (BL –) or blue light (BL +) at 10 µmol m^−2^ s^−1^ for 16 min in the presence of γ-^32^P-ATP. The reacted samples were resolved on 6% SDS-PAGE gels and autoradiographed. **(B)** Relative quantify of phosphorylated 120-kDa proteins. The value was calculated against the data from Est-untreated, blue light-irradiated *pMDC7-RPT2* seedlings. The data shown are mean values ± SE (*n* = 3). An asterisk indicate a statistically significant difference (Student’s t test; *P* < 0.05).

Our present analyses suggested that RPT2 proteins either suppress phot1 autophosphorylation or enhance its dephosphorylation. If a loss-of-function mutation in *RPT2* leads to an enhanced phosphorylation of phot1 due to a block in its dephosphorylation and RPT2 overexpression enhances its dephosphorylation, treatments with protein phosphatase inhibitors seemed to impair the effects of a loss-of-function mutation and overexpression of *RPT2*. We therefore next examined the effects of protein phosphatase inhibitors in the *rpt2* mutants and the *35Spro:RPT2* transgenic lines by immunoblotting analysis using a Phos-tag acrylamide gel. First, we examined the effects of the protein phosphatase inhibitors cantharidin (CN) or okadaic acid (OKA) on the dephosphorylation of phot1. Autophosphorylated PHOT1 proteins in the wild-type seedlings with a pulse irradiation of blue light for 2 min at 100 µmol m^−2^ s^−1^ were dephosphorylated with a subsequent dark incubation for 14 min (Supplemental Figure 7). When the seedlings were treated with CN or OKA, the mobility of the PHOT1 protein became slightly retarded in Phos-tag SDS-PAGE (Supplemental Figure 7). These results indicated that CN and OKA have some inhibitory effects on phot1 dephosphorylation.

We next examined the effects of these inhibitors in the *rpt2* mutants. When wild-type seedlings were irradiated with blue light for 0.5 h in the CN- or OKA-containing medium with a red-light pretreatment (for the induction of RPT2: Haga et al. 2015), the PHOT1 proteins showed a hyper mobility shift in contrast to the untreated seedlings (Figure 4A). The phosphorylation of phot1 was enhanced in the *rpt2* mutants in comparison with those in wild-type seedlings independently of the treatments of CN or OKA (Figure 4A). The effects of CN and OKA in the *35Spro:RPT2* transgenic line were also examined (Figure 4B). Red light pretreatment was not done here to ensure that RPT2 expression was not induced in the wild-type seedlings. Constitutive expression of RPT2 suppressed the phosphorylation of phot1 in the *35Spro:RPT2* transgenic lines without being affected by the CN and OKA treatments (Figure 4B). These results suggested that RPT2 suppresses the autophosphorylation activity of phot1 but does not enhance the dephosphorylation of phot1.

**Figure 4.**
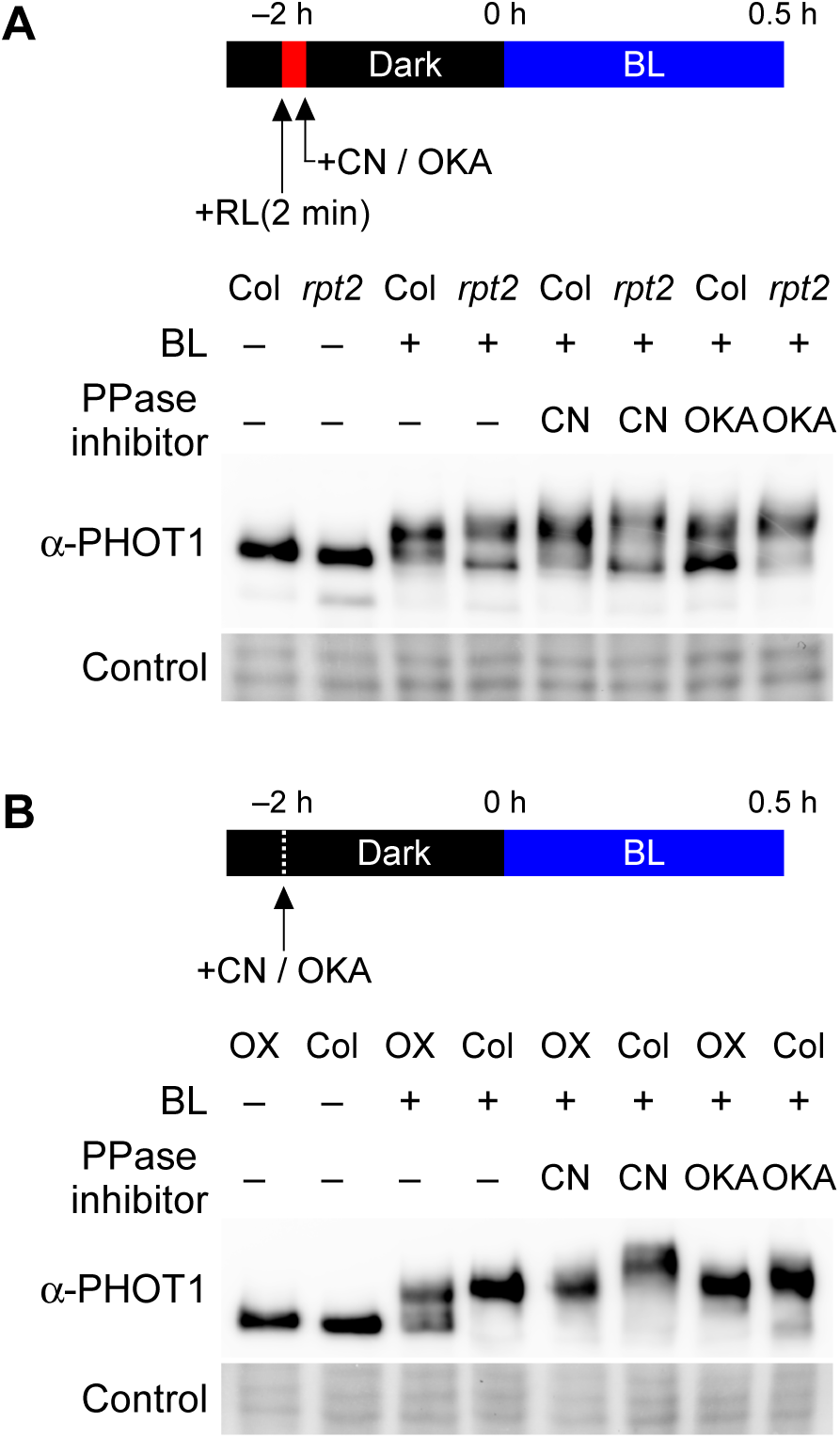
Effect of protein phosphatase inhibitors on phosphorylation status of phot1. The aerial parts of two-day-old etiolated seedlings were incubated under darkness for 2 h in liquid medium with a protein phosphatase inhibitor (PPase inhibitor), 30 µM cantharidin (CN) or 1 µM okadaic acid (OKA), and subsequently treated with blue light at 0.1 µmol m^−2^ s^−1^ for 0.5 h. Total proteins (15 µg for **A**, 20 µg for **B**) extracted from the seedlings were separated on 6% SDS-PAGE gels with 2 µM Phos-tag, followed by immunoblotting with α-PHOT1 antibodies. Protein-blotted membranes were stained as a loading control. **(A)** Effect of PPase inhibitors on the aerial parts of wild-type (Col) seedlings and *rpt2* mutants. Red light pretreatments were conducted to induce RPT2 expression before exposure to the PPase inhibitors. **(B)** Effect of PPase inhibitors on the aerial parts of wild-type (Col) seedlings and the *35Spro:RPT2* lines (OX).

### RPT2 is induced by blue-light irradiation in a post-transcriptional manner

Our current results suggested that the induction of RPT2 expression by light irradiation is an important mechanism for controlling both photosensitivity and the autophosphorylation activity of phot1. We thus investigated the light inducibility of RPT2 expression in more detail. We first observed the RPT2 expression patterns using transgenic plants carrying the *RPT2pro:GUS* gene and the *RPT2pro:RPT2-VENUS* gene. GUS staining was detected in a root tip under darkness in the 2-day-old etiolated seedlings carrying the *RPT2pro:GUS* gene (Figure 5A, 5D, and 5G). Both red- and blue-light irradiation enhanced its expression in whole seedlings, most notably the hypocotyls, hooks, and root tips including the elongation zone, in a similar manner (Figure 5B, 5C, 5E, 5F, 5H, and 5I). When the expression patterns of the *RPT2pro:RPT2-VENUS* gene in the *rpt2* mutants were analyzed, we noticed a clear induction of RPT2-VENUS proteins in the aerial part of seedlings (Figure 5L and 5S), especially the elongation zones of the hypocotyls (Figure 5O), and the elongation zone of roots (Figure 5R), but only under blue-light irradiation and not red-light irradiation (Figure 5K, 5N and 5Q). These results indicated that both red- and blue-light irradiation can activate the *RPT2* promoter but that only blue-light irradiation can induce the RPT2-VENUS proteins effectively.

**Figure 5.**
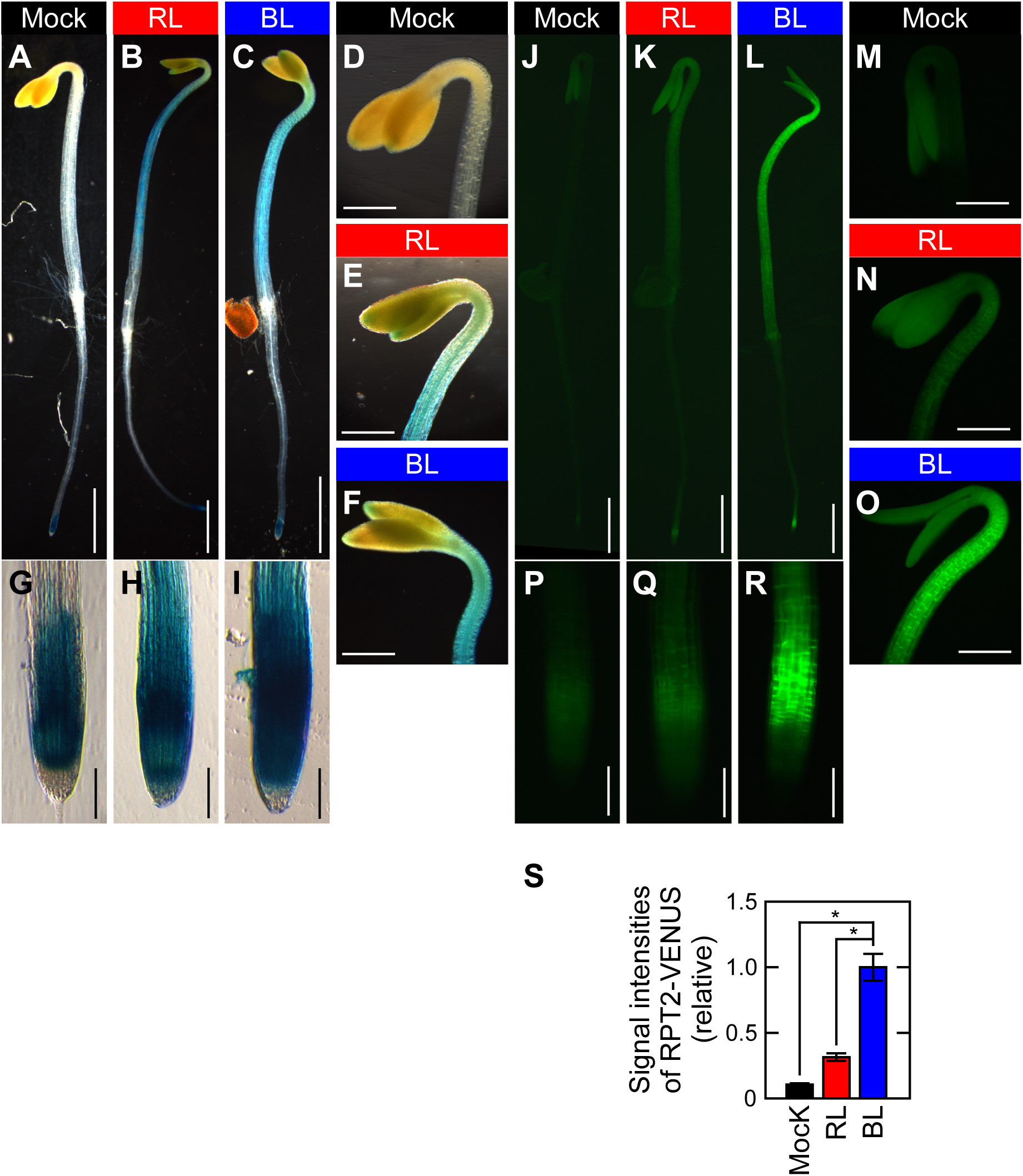
Expression patterns of the *RPT2pro:GUS* and the *RPT2pro:RPT2-VENUS* genes. Two-day-old etiolated seedlings were mock-irradiated (Mock: **A, D, G, J, M, P**), or red light-(RL: **B, E, H, K, N, Q**) or blue light-(BL: **C, F, I, L, O, R**) irradiated at 10 µmol m–2 s–1 for 4 h. Scale bars: 1.0 mm **(A-C, J-L)**; 400 µm **(D-F, M-O)**; 100 µm **(G-I, P-R)**. **(A-I)** GUS staining patterns of wild-type seedlings transformed with a *RPT2pro:GUS* gene. **(J-R)** VENUS fluorescent images of the *rpt2* mutants transformed with a *RPT2pro:RPT2-VENUS* gene. **(S)** Signal intensities of RPT2-VENUS fluorescence. Fluorescent signals were measured in the upper region of the hypocotyls and calculated relative to the value from BL-irradiated seedlings. The data shown are the mean values ± SE from 9 seedlings. Asterisks indicate a statistically significant difference (Student’s t test; *P* < 0.01).

### The accumulation of RPT2 proteins is enhanced by phot activation

The RPT2-VENUS fluorescent signal was detectable in the elongation zones of hypocotyls and roots (Figure 5O and 5R), in which the strong expression of phot1 have been reported (Sakamoto and Briggs, 2002). Thus, we hypothesized that RPT2 proteins are unstable under red-light conditions and that phot1 stabilizes them under blue-light conditions. We tested this using immunoblotting analysis. When the wild-type seedlings were irradiated for 6 hours with red or blue light at 10 µmol m^−2^ s^−1^, RPT2 protein expression was induced in both cases but blue-light irradiation was much more effective (Figure 6A and 6B). On the other hand, the accumulation of RPT2 proteins under blue-light conditions was attenuated in the *phot1* mutants, which was statistically significant (Figure 6A and 6B). The mutation of the *NPH3* gene, which is required for the phot1 signaling during phototropism (Motchoulski and Liscum, 1999), also caused a decrease of RPT2 accumulation under blue-light conditions (Figure 6A and 6B). Neither *phot1* nor *nph3* mutations affected the red-light induced accumulation of RPT2 (Figure 6A and 6B). qRT-PCR analysis confirmed that the blue-light induction of *RPT2* transcripts was not affected by mutations of *phot1* and *nph3* (Figure 6C). These results suggested that phot1 and its associated protein NPH3 contribute to the accumulation of RPT2 proteins under blue-light conditions in a post-transcriptional manner.

**Figure 6.**
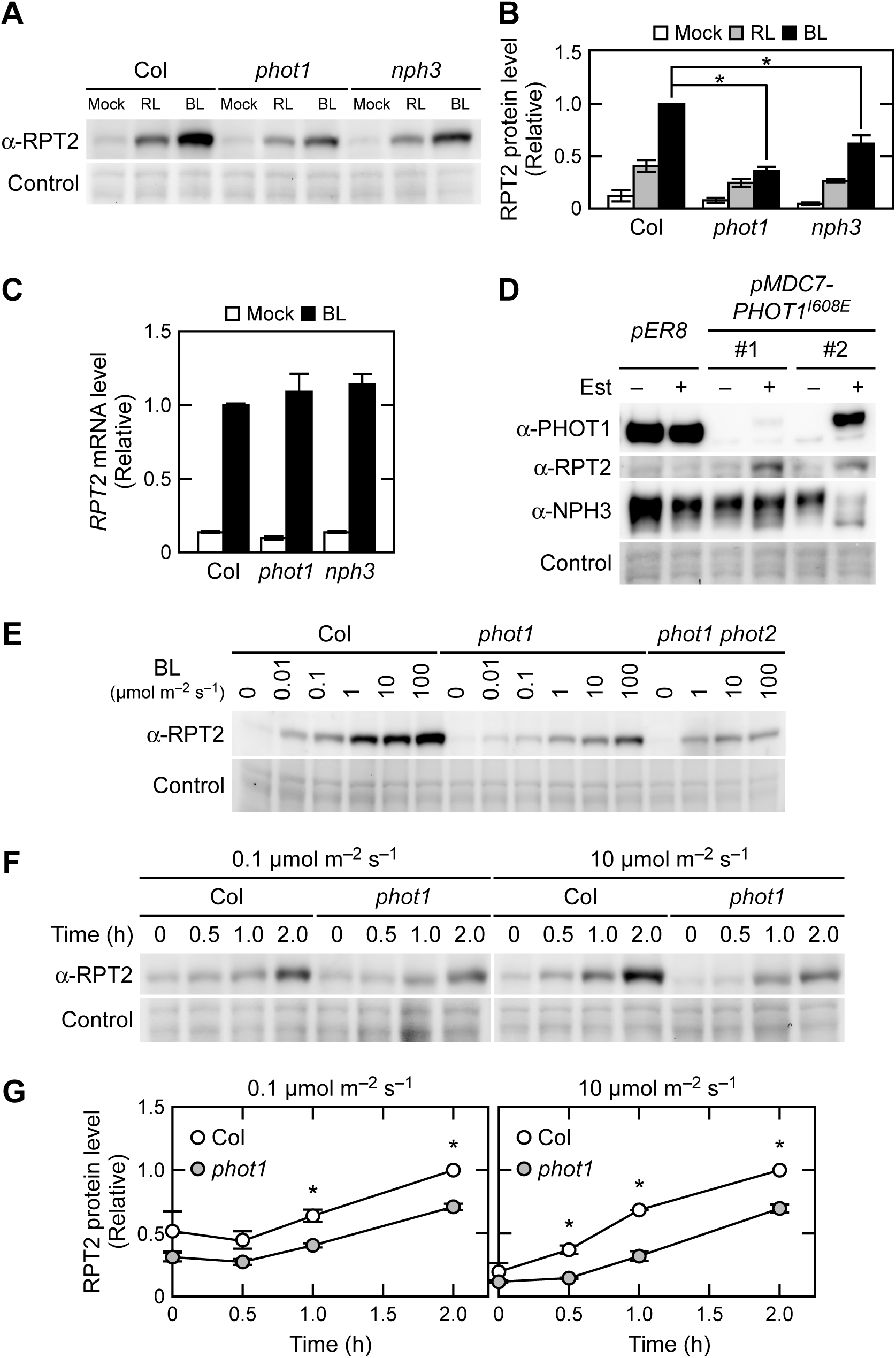
Post-transcriptinal regulation of RPT2 expression by phototropins. **(A)** Immunoblotting analysis of RPT2 proteins in wild type (Col), *phot1*, and *nph3* seedlings. Two-day-old etiolated seedlings were mock-irradiated (Mock), or red light-(RL) or blue light-(BL) irradiated at 10 µmol m^−2^ s^−1^ for 6 h. Total proteins (10 µg) extracted from the seedlings were resolved on 10% SDS-PAGE gels, followed by immunoblotting with α-RPT2 antibodies. The protein-blotted membranes were stained as a loading control. **(B)** Statistical analysis of the data in **(A)**. The values were normalized with a loading control and then calculated against the data from the blue light-irradiated seedlings of wild type. The data shown are the mean values ± SE (*n* = 3). Asterisks indicate a statistically significant difference (Student’s t test; *P* < 0.05). **(C)** qRT-PCR analysis of *RPT2* in wild type (Col), *phot1*, and *nph3* seedlings. Two-day-old etiolated seedlings were irradiated with blue light at 10 µmol m^−2^ s^−1^ for 6 h or mock-irradiated. The values were normalized using an internal control (*18S rRNA*) and the calculated against the values from blue light-irradiated seedlings of wild type. The data represent the means and SE (*n* = 3). **(D)** Immunoblotting analysis of PHOT1, RPT2 and NPH3 in phot1 mutants transformed with a *pMDC7-PHOT1^I608E^* construct (*pMDC7*-*PHOT1^I608E^*: two independent lines, #1 and #2) and the pER8 vector control line. Two-day-old etiolated seedlings were grown on agar medium with (+) or without (–) 10 µM estradiol (Est). Microsomal proteins (7.5 µg) extracted from the seedlings were separated on 7.5% SDS-PAGE gels, followed by immunoblotting with α-PHOT1, α-RPT2, and anti-NPH3 (α-NPH3) antibodies. The protein-blotted membranes were stained as a loading control. **(E)** Fluence-rate dependency of the RPT2 induction in wild type (Col), *phot1*, and *phot1 phot2* double mutants. Two-day-old etiolated seedlings were irradiated with blue light at the indicated fluence rate for 6 h. Other details were as described in **(A)**. **(F)** Time course analysis of RPT2 induction in wild type (Col) and *phot1* seedlings. Two-day-old etiolated seedlings were irradiated with blue light at 0.1 (left panel) or 10 µmol m^−2^ s^−1^ (right panel) for the indicated period. Other details were as described in **(A)**. **(G)** Statistical analysis the data generated in **(F)**. The values were normalized using a loading control and then calculated against the values from wild-type seedlings irradiated for 2 h. The data shown are the mean values ± SE (*n* = 3). Asterisks indicate a statistically significant difference (Student’s t test; *P* < 0.05).

We next examined whether the sole activation of the phot1 photoreceptor is sufficient for the accumulation of RPT2 proteins using *phot1* mutants transformed with a *pMDC7-PHOT1^I608E^* construct, in which constitutively active PHOT1 proteins (PHOT1^I608E^: Harper et al., 2004) are inducible by Est treatment. In the etiolated seedlings of the *pER8* vector control line, Est treatment had no effect on the RPT2 protein levels or the phosphorylation status of the NPH3 protein (Figure 6D). On the other hand, in the *phot1* mutants transformed with a *pMDC7-PHOT1^I608E^* construct (#1 and #2), Est exposure caused the accumulation of RPT2 protein in the absence of blue-light irradiation. This suggested that the accumulation of RPT2 can be caused by phot1 activation alone, even in darkness.

The mobility shift of the NPH3 protein, which reflects a phot1-induced dephosphorylation (Pedmale and Liscum, 2007; Tsuchida-Mayama et al., 2008), was also observed in the #2 transgenic line (Figure 6D), suggesting that *phot1^I608E^* expression can cause NPH3 dephosphorylation under darkness. A previous study reported that the expression of *PHOT1^R472H^*, which is another constitutively active variant of phot1, was not sufficient to promote NPH3 dephosphorylation under darkness (Petersen et al., 2017). The discrepancies between the prior result and our current observations may be due to differences in the natures of phot1^I608E^ and phot1^R472H^ and/or the transgene expression levels.

We next analyzed the fluence-rate and time dependence of RPT2 protein accumulation. When wild-type seedlings were irradiated with blue light at 0.01 to 100 µmol m^−2^ s^−1^ for 6 hours, the RPT2 protein levels were increased as the fluence rates of blue light increased (Figure 6E). This induction seemed to be caused by both a transcriptional regulation of phy and cry (Tsuchida-Mayama et al., 2010) and a post-transcriptional regulation of phot1. In the *phot1* mutants, a weakened induction of the RPT2 protein was observed at all fluence rates examined (Figure 6E), suggesting that phot1 contributes to RPT2 accumulation, at least between 0.01 to 100 µmol m^−2^ s^−1^. In the *phot1 phot2* double mutants, the blue-light induction of RPT2 at 100 µmol m^−2^ s^−1^ was obviously lower than that in the *phot1* single mutant (Figure 6E), suggesting that phot2 also contributes to the accumulation of RPT2 proteins at 100 µmol m^−2^ s^−1^. When the seedlings were irradiated by blue light at 0.1 or 10 µmol m^−2^ s^−1^ for various times, the induction of RPT2 proteins was detectable at least after 1 hour of the onset of irradiation in both wild type and *phot1* seedlings, but at a higher level in wild type (Figure 6F and 6G). This observation suggested that both the transcriptional induction of *RPT2* by phys and crys (Tsuchida-Mayama et al., 2010) and the post-transcriptional induction of phot1 contributes to an early induction of RPT2 under both fluence-rate conditions.

Previous study has indicated that phot1 is localized to the plasma membrane region in the epidermal cells and the cortical cells of both root and hypocotyl of Arabidopsis etiolated seedlings (Sakamoto and Briggs, 2002). When the *rpt2* transgenic seedlings expressing RPT2-VENUS were irradiated by blue light from above, a fluorescent image of a transverse section of upper hypocotyls showed that the RPT2-VENUS proteins were localized to the plasma membrane region in all tissues of hypocotyls and were strongly expressed in the cortex (Figure 7A). These expression and subcellular localization patterns were similar with those of phot1 (Sakamoto and Briggs, 2002).

**Figure 7.**
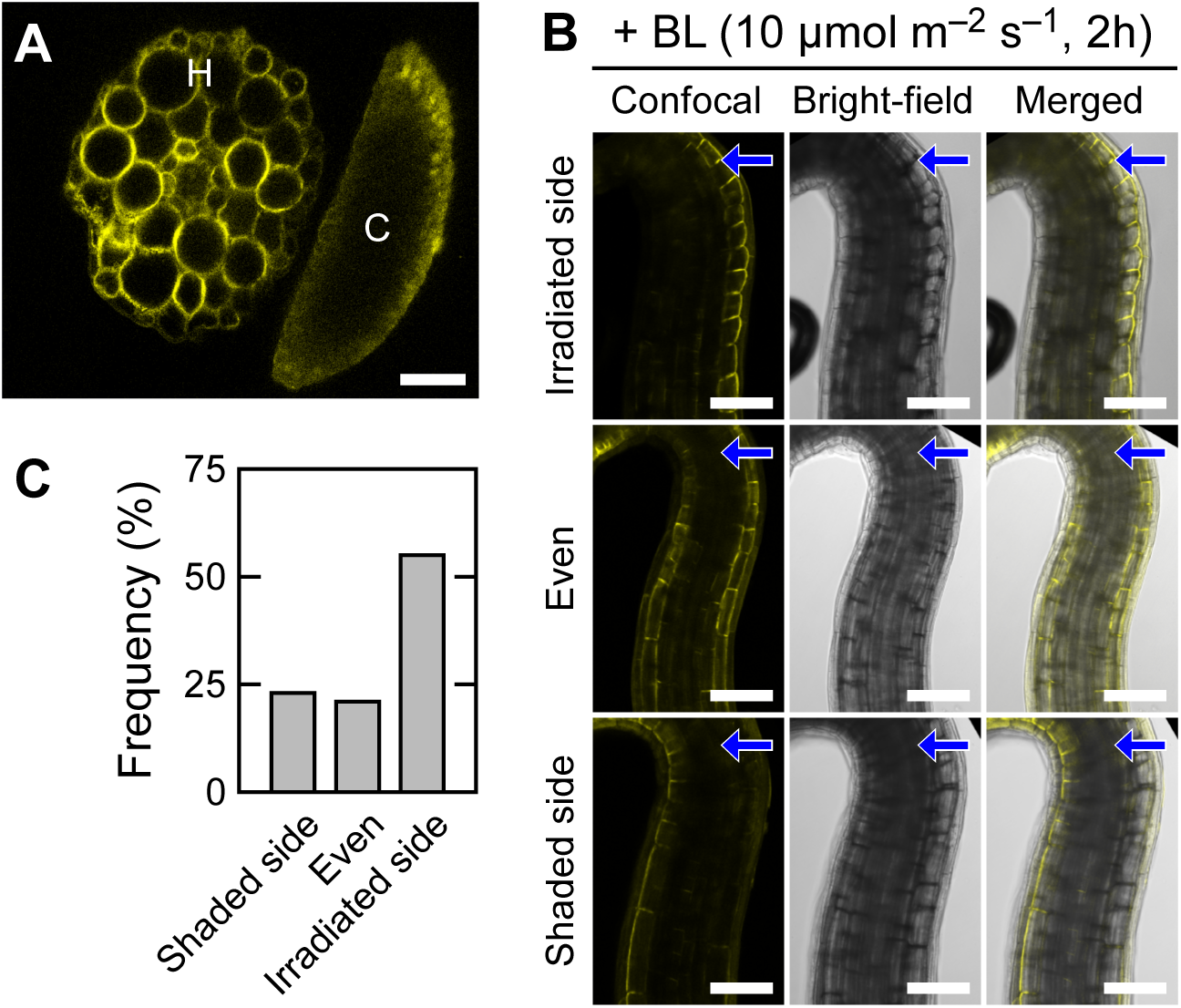
Distribution pattern of RPT2-VENUS protein in etiolated hypocotyls. Two-day-old etiolated seedlings of *rpt2* mutants transformed with a *RPT2pro:RPT2-VENUS* gene were irradiated with blue light (BL) at 10 µmol m^−2^ s^−1^ for 6 h from above **(A)** or for 2 h from the unilateral side **(B)**. **(A)** A typical distribution patten of RPT2-VENUS in the hypocotyl cross section. H, hypocotyl; C, cotyledon. **(B and C)** Distribution patterns of RPT2-VENUS in the upper region of the hypocotyls. Distribution patterns were classified into three types (Irradiated side, Even and Shaded side). Representative confocal (left panel), bright field (mid panel) and merged images (right panel) are shown in **(B)**. Arrows indicate the direction of blue light. The frequencies of each expression pattern were calculated from 74 seedlings and are shown in **(C)**. White bar, 100 µm.

Suzuki et al. (2019) has reported that unilateral irradiation of blue light induces the asymmetric distribution of phosphorylated Zmphot1 in coleoptiles of *Zea mays* in response to the gradient of blue light intensity in these organs. When the seedlings were unilaterally irradiated, the RPT2-VENUS proteins were often expressed more strongly on the irradiated side of the hypocotyls than on the shaded side (∼55% of seedlings; Figure 7B and 7C). On the other hand, there were also seedlings showing symmetric expression patterns (∼21%) or a strong expression of RPT2-VENUS on the shaded side (∼24%). As the light induction of endogenous RPT2 proteins was already detectable at 1 h after the onset of blue light irradiation at 10 µmol m^−2^ s^−1^ in the immunoblotting analysis (Figure 6E), their distribution patterns at 1 h could also be observed. The RPT2-VENUS fluorescent signal, however, was barely detectable (data not shown). One hour of irradiation seemed to be too short for the expression, maturation and accumulation of RPT2-VENUS proteins to detect its fluorescence. Therefore, we could not draw any conclusions regarding the asymmetric induction of RPT2 in hypocotyls irradiated by unilateral blue light in our present study.

### RPT2 proteins are degraded through the ubiquitin-proteasome pathway

We examined whether the RPT2 protein is degraded through a ubiquitin-proteasome pathway. When etiolated seedlings of wild type and *phot1 phot2* double mutants were treated with the proteasome inhibitor MG132 for 3 hours under blue light conditions at 10 µmol m^−2^ s^−1^, RPT2 protein accumulated in the double mutant but not in wild type. This suggested that the *phot1 phot2* double mutants showed a destabilization of RPT2 proteins in a proteasome-dependent manner (Figure 8A). In addition, exposure to MG132 led to the accumulation of RPT2 in wild-type seedlings under red-light conditions but not under blue-light conditions (Supplemental Figure 8). These results suggested that RPT2 is degraded in a proteasome-dependent manner and is stabilized though the activation of phototropins.

**Figure 8.**
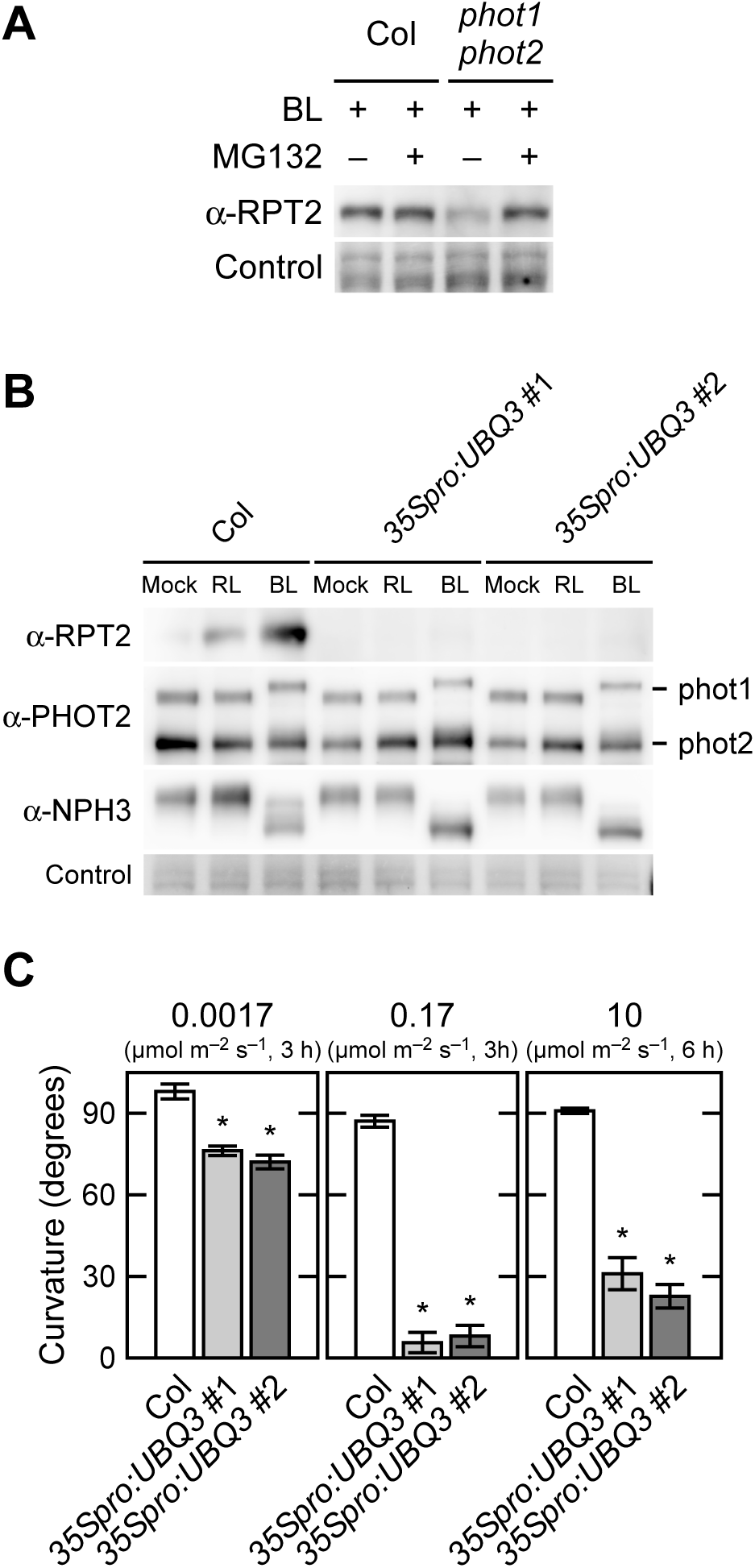
Destabilization of the RPT2 protein via a ubiquitin-proteasome dependent pathway. **(A)** Effect of the proteasome inhibitor MG132 on RPT2 protein expression. Two-day-old etiolated seedlings of wild type (Col) and *phot1 phot2* double mutant were transferred into liquid medium with (+) or without (–) 50 µM MG132, kept in the dark for 1 h, and subsequently irradiated with blue light (BL) at 10 µmol m^−2^ s^−1^ for 3 h (+) or mock-irradiated (–). Total proteins (10 µg) extracted from the seedlings were separated on 10% SDS-PAGE gels, followed by immunoblotting with α-RPT2 antibodies. The protein-blotted membranes were stained as a loading control. **(B)** Immunoblotting analysis of RPT2, PHOT1, PHOT2 and NPH3 in wild-type (Col) and wild-type seedlings transformed with *35Spro:UBQ3 (35Spro:UBQ3*: two independent lines; #1 and #2). Two-day-old etiolated seedlings were mock-, red light-(RL) or blue light-(BL) irradiated at 10 µmol m^−2^ s^−1^ for 6 h. Total proteins (10 µg) extracted from the seedlings were then separated on 7.5% SDS-PAGE gels, followed by immunoblotting with α-RPT2, α-PHOT2 and α-NPH3 antibodies. The protein-blotted membranes were stained as a loading control. **(C)** Hypocotyl phototropism in wild-type (Col) and *35Spro:UBQ3* transgenic lines. Two-day-old etiolated seedlings were irradiated with unilateral blue light at 0.0017 or 0.17 µmol m^−2^ s^−1^ for 3 h, or 10 µmol m^−2^ s^−1^ for 6 h. The data shown are the mean values ± SE for hypocotyl curvatures of 14-24 seedlings. Asterisks indicate a statistically significant difference (Student’s t test; *P* < 0.05).

To detect polyubiquitinated RPT2 under red-light irradiation, we attempted to immunoprecipitate it from extracts of *UBIQUITIN3* (*UBQ3*) overexpression lines (*35Spro:UBQ3*). Although this was not successful, we unexpectedly found a significant decrease of RPT2 protein in the *35Spro:UBQ3* transgenic lines under both red- and blue-light conditions (Figure 8B). Correspondingly, these transgenic lines showed an abnormality in hypocotyl phototropism i.e. their phototropic curvatures were moderate under weak intensity blue-light conditions and decreased as the fluence rate increased (Figure 8C), in a comparable manner to the *rpt2* mutants (Haga et al., 2015). On the other hand, *UBQ3* overexpression had no effect on the expression patterns of PHOT1, PHOT2 or NPH3 (Figure 8B), or on the growth of seedlings (Supplemental Figure 9A and 9B). These results suggested that the RPT2 proteins are degraded through the ubiquitin-proteasome pathway and that the activation of phot1 and phot2 may negatively regulate the polyubiquitination of RPT2 proteins and/or its degradation by proteasomes.

## DISCUSSION

Our current results demonstrate that RPT2 suppresses the autophosphorylation of phot1 under blue-light conditions. Previous studies have indicated that the phot1 LOV1 domain is necessary for the induction of the phototropic responses under low intensity blue-light conditions (Sullivan et al., 2008) and that the light induction of RPT2 expression suppresses phot1 photosensitivity, which is required for the photosensory adaptation of phot1 to high intensity blue light (Haga et al., 2015). These prior results and our current data suggest that RPT2 negatively regulates the autophosphorylation of phot1 through the LOV1 domain, which is probably required for a formation of a suitable gradient of phot1 activity between the irradiated side and the shaded side in accordance with the high intensity unilateral blue light. Hence, this is the photosensory adaptation mechanism of phot1 in the second positive phototropism of Arabidopsis etiolated hypocotyls. The expression level of RPT2 proteins is increased in response to an increase of blue light intensity (Figure 6E), indicating that RPT2 functions as a molecular rheostat that maintains a moderate activation level of phot1 in etiolated hypocotyls of *Arabidopsis* seedlings under any light intensity conditions.

The molecular mechanisms by which RPT2 functions in the suppression of phot1 activity remain to be elucidated. Some prior studies have suggested that the LOV1 domain mediates the dimerization of phot1, which thereby enhances the autophosphorylation of this blue-light photoreceptor (Nakasako et al., 2004; Xue et al., 2018). Thus, RPT2 may inhibit the LOV1-mediated dimerization of phot1 to suppress its kinase activity. Previous studies had also suggested that LOV1 suppresses the decay of the cysteinyl-FMN adduct of LOV2 and enhances the Ser/Thr kinase activity of the phots (Kaiserli et al., 2009; Okajima et al., 2012; Okajima 2016). RPT2 may enhance the decay of the cysteinyl-FMN adduct of phot1 LOV2 through the binding to LOV1 and thus suppress the Ser/Thr kinase activity of phot1. Although our current results suggest that RPT2 suppresses the autophosphorylation activity of phot1, the possibility of enhancement of phot1 dephosphorylation by RPT2 still cannot be excluded. RPT2 might play a role as a scaffold of protein phosphatases to dephosphorylate phot1. We also need to examine in the future whether a defect of RPT2 binding to the LOV1 domain indeed affects the autophosphorylation of phot1. Further studies are warranted to elucidate the mechanisms underlying the function of the LOV1 domain and RPT2 in phototropin photoactivation in more detail.

The post-transcriptional regulation of RPT2 forms a negative feedback loop of phot1 activation. This regulation appears to ensure the formation of a gradient of phot1 signaling activity between the irradiated and shaded sides of plant organs under a broad range of blue-light intensity. This gradient then seems to induce light-induced differential growth including not only phototropic responses but also leaf flattening and cotyledon/leaf positioning (Sakai et al., 2000; Harada et al., 2013). Previous studies have reported that unilateral blue light irradiation results in differential NPH3 aggregate formation in response to phot1 activity across the etiolated hypocotyl, which suppresses and fine-tunes NPH3 activity (Sullivan et al., 2019), and that the light-induced RPT2 proteins suppresses the dephosphorylation and aggregation of NPH3 proteins in the etiolated seedlings (Haga et al., 2015). Our current study findings strongly suggest that RPT2 indirectly suppresses their dephosphorylation and aggregation through the suppression of phot1 activity. On the other hand, our current study results also indicate that NPH3 partially contributes to the stabilization of RPT2 proteins under blue-light conditions (Figure 6B). Thus, the aggregation of NPH3 proteins might decrease the stabilization of RPT2 proteins at the irradiated side and fine-tune the phot1 activity at both the irradiated and the shaded side. Both adjustments of the light induction of PRT2 and of the plasma membrane localization of NPH3 probably contribute to the fine-tuning of phot1 signaling across hypocotyls, and an induction of phototropic responses under various light conditions in the etiolated seedlings of *Arabidopsis*. On the other hand, RPT2 is not required for the second positive phototropism in de-etiolated hypocotyls of Arabidopsis seedlings (Sullivan et al., 2019). Other unknown factors and/or mechanisms may suppress the excessive activation of phot1 in green seedlings.

Our current observations indicate that RPT2 is degraded by the ubiquitin-proteasome pathway and that phot1 activation suppresses this degradation. This is the first demonstration of a protein stabilization control function of the phots. The mechanism by which the RPT2 protein is ubiquitinated or stabilized under blue-light conditions has been a question of some importance. NPH3 is an essential signal transducer during phototropism and shows binding activity towards RPT2 and possesses ubiquitin E3 ligase activity with Cullin3 (Motchoulski and Liscum, 1999; Inada et al., 2004; Roberts et al., 2011). Although we speculated that NPH3 may ubiquitinate RPT2 and promote its degradation under red-light conditions, our immunoblotting analysis revealed that NPH3 contributes to its accumulation under blue-light conditions (Figure 6A). As RPT2 belongs to the same protein family as NPH3 and also has a BTB/POZ domain which often interacts with Cullin3 (Genschik et al., 2013), it may be ubiquitinated on its own, and its binding to the active forms of the phots may suppress its polyubiquitination and degradation. The issue of whether phot1 controls a ubiquitin-proteasome pathway also remains to be resolved.

*RPT2* belongs to the *NPH3/RPT2-like* (*NRL*) gene family (Sakai, 2005) and other NRL members might also have a similar function to RPT2. RPT2 did not show an obvious effect on the suppression of the phot2 activity, although it can bind to its LOV1 domain and form a complex with phot2 in vivo. However, for example, NRL PROTEIN FOR CHLOROPLASTMOVEMENT1 (NCH1) functions in the chloroplast accumulation response in parallel with RPT2 (Suetsugu et al., 2016). Suetsugu et al. (2016) revealed that *nch1* mutants show an enhancement of the phot2-induced avoidance response. One of the hypotheses from this is that NCH1 suppresses the photosensitivity and/or photoactivation of phot2 with RPT2. Thus, the relationships between the NRL members and phototropins in various plants should be reexamined in future studies.

## METHODS

### Plant Materials and Growth Conditions

*Arabidopsis thaliana* ecotype Columbia (Col) was used as the wild type control. Mutant seeds of *rpt2-2* (Col background) and *nph3-102* (Salk_110039; Col background) were obtained as described previously (Inada et al., 2004; Tsuchida-Mayama et al., 2008). The *phot1-*Salk146058 (Col background) and *phot2*-Salk142275 mutants (Col background) were obtained from the Arabidopsis Biological Resource Center (Alonso et al., 2003) and crossed to obtain a *phot1 phot2* double mutant. The *rpt2-2* mutants transformed with a *35Spro:RPT2* gene or a *RPT2pro:RPT2-VENUS* gene were prepared as described previously (Tsuchida-Mayama et al., 2010; Haga et al., 2015). Transgenic Col lines harboring a *RPT2pro:GUS* gene were also prepared as described previously (Inada et al., 2004). The *phot1 phot2* mutants transformed with a *35Spro:PHOT1^LOV1Cys39Ala^* gene were kindly provided by Professor John Christie (University of Glasgow).

The *pMDC7-RPT2* transgenic lines were generated as follows. The *RPT2* coding region was subcloned into the entry vector pENTR/D-TOPO (Invitrogen) and shuttled via an LR clonase reaction (Invitrogen) into the estrogen (Est)-inducible vector pMDC7 (Curtis and Grossniklaus, 2003), which was kindly provided by Professor Nam-Hai Chua (Rockefeller University, New York, NY). The *pMDC7-RPT2* plasmid was used for the transformation of *Agrobacterium tumefaciens*, which was then used for subsequent transformations of *rpt2-2* mutants. Several independent *rpt2* transgenic lines showed phototropism complementation following Est treatment (Supplemental Figure 6A). pER8 is an original vector of pMDC7 that lacks the Gateway cassette (Zuo et al., 2000) and was used for the transformation of *A. tumefaciens*, which was then used for subsequent transformations of Col wild type for use as a vector control line.

The *pMDC7-PHOT1^I608E^* transgenic lines were generated as follows. The *PHOT1^I608E^* mutated cDNA was generated from *PHOT1* cDNA by PCR-based site-directed mutagenesis using *PHOT1*-gene specific primers including 5’-A<u>CTGCAG</u>TTTTTTCACCAGGTCTTCTCCCTC-3’ and 5’-A<u>CTGCAG</u>TGAAT<u>GAA</u>GATGAAGCGGTTCGAGAACT-3’ (underlined bases denote the Glu [E] codon 608; double underlines indicate the PstI sites for subsequent subcloning), subcloned into the entry vector pDONR222 (Invitrogen), and shuttled into pMDC7 via an LR clonase reaction. The *pMDC7-PHOT1^I608E^* plasmid was used for the transformation of *A. tumefaciens*, which was then used for subsequent transformations of *phot1*-Salk146058 mutants.

The *35Spro:UBQ3* transgenic lines were generated as follows. The POLYUBIQUITIN 3 (UBQ3) coding region was amplified from Col genome DNA by PCR using *UBQ3*-gene specific primers (Supplemental Table S1), subcloned into the entry vector pENTR/D-TOPO, and shuttled via an LR clonase reaction into the cauliflower mosaic virus 35S promoter-containing binary vector pH35GS, which was kindly provided by Professor Taku Demura (Nara Institute of Science and Technology, Nara, Japan). The resulting *pH35GS-UBQ3* plasmid was used for the transformation of *A. tumefaciens*, which was then used for subsequent transformations of wild type (Col) plants.

For experiments, the seeds were surface-sterilized and plated in Petri dishes with half-strength Okada and Shimura medium containing 1.5% agar, as described previously (Ohgishi et al., 2004). Seeds were kept at 4°C for 3 days and then exposed to red light for 6 h to induce uniform germination. After germination was induced, the Petri dishes were positioned vertically to let the seedlings grow on the surface of the agar at 21 to 22°C under dark conditions. Blue-light and red-light irradiations were performed under various conditions with light-emitting diodes (Ohgishi et al., 2004), as described in the Figure legends.

### Yeast Two-Hybrid Analysis

The GAL4 DNA-binding domain vector pGBDKT7-GWRFC was constructed by insertion of the Gateway reading frame cassette RfcC (Invitrogen) into the Klenow-Fragment-treated NdeI-SalI sites of pGBKT7 (pGBDKT7: Clontech, http://www.clontech.com). PCR was used to generate the coding sequences of PHOT1 and PHOT2 with gene specific primers (Supplemental Table S1). A DNA fragment for PHOT1 N2mut was separately generated by PCR using N2 FW and N2mut RV and N2mut FW and N2 RV primers, and then combined by PCR using N2 FW and N2 RV primers. These amplified products were subcloned into the entry vector pENTR/D-TOPO and shuttled into the pGBDKT7-GWRFC plasmid. The GAL4 transcription-activating domain vector pGADT7 and its derivative pGADT7-RPT2 N were prepared as described previously (Inada et al., 2004). Pairwise combinations of vectors were co-transformed into the yeast strain Y187 (Clontech) and plated onto the same selective medium. Quantitative β-galactosidase assays were performed in liquid cultures of yeast using *o*-nitrophenyl-β-D-galactopyranoside (Ausubel et al., 2001). One unit of β-Gal activity was defined as the amount of enzyme required to convert 1 µmol of *o*-nitrophenyl-β-D-galactopyranoside to *o*-nitrophenol and D-galactose in 1 min at 30°C.

### In vitro Pull-Down Assay

To prepare phot1 LOV1 and LOV2 proteins, DNA fragments for *PHOT1 N2* and *N4* in pENTR/D-TOPO (Invitrogen) were transferred into the pCold ProS2 plasmid (TaKaRa) harboring a gateway reading flame cassette (Invitrogen). These constructs were introduced into *Escherichia coli* strain BL21 (DE3) pLysS (Novagen), and His-ProS-tagged PHOT1 N2 and N4 proteins (His-PHOT1N2 and –PHOT1N4) were prepared from the transformed lines in accordance with the manufacturer’s protocol. HA-tagged RPT2 N proteins were prepared with *in vitro* transcription and translation of pGADT7-RPT2 N using the TNT Quick Coupled Transcription/Translation Systems (Promega).

Each purified protein preparation of His-PHOT1 N2 and N4 was incubated with TALON Magnetic Beads (TaKaRa) at 4°C for 30 min and further incubated at 4°C for 30 min with *in vitro* transcription and translation reactant containing RPT2 N. The beads were then collected on the magnetic rack and washed five times with washing buffer (sodium phosphate buffer pH 7.0, 150 mM NaCl, 0.2% Triton-X100). The proteins were then released from the beads into 50 µL of 1× SDS gel loading buffer and resolved by SDS-PAGE.

### Immunoblotting Analysis

For immunoblotting of Phos-tag SDS-PAGE gels, total proteins were extracted from etiolated seedlings in a buffer containing 50 mM Tris-MES, pH 7.5, 300 mM sucrose, 150 mM NaCl, 10 mM potassium acetate, 0.2% Triton X-100, and a protease inhibitor mixture (Complete Mini EDTA-free; Roche Diagnostics). The extracts were then centrifuged at 10,000 g at 4°C for 10 min to remove cell debris and the supernatants were collected and mixed with a half volume of 3× SDS gel loading buffer, and boiled at 95°C for 15 min. The samples were separated with 6% SDS-PAGE gels containing 2 µM Phos-tag (FUJIFILM Wako Pure Chemical Corporation) in accordance with the previously described “Zn^2+^-Phos-tag SDS PAGE” method (Kinoshita and Kinoshita-Kikuta, 2011). Following electrophoresis, the gels were twice washed with methanol-free transfer buffer (25 mM Tris, 192 mM glycine) with 10 mM EDTA for 10 min and then once with methanol-free transfer buffer without EDTA for 10 min. The separated total proteins were then blotted onto a PVDF membrane using a wet tank blotting system at a constant voltage of 350 mA for 4 h. Later immunoblotting steps were performed as described previously (Inada et al., 2004). For alkaline phosphatase treatment, microsomal pellets were obtained as described previously (Inada et al., 2004) and resuspended in the 1× NEBuffer 3 (NEB) with 0.5% Triton-X100 and protease inhibitor mixture (Complete Mini EDTA-free; Roche Diagnostics). 30 µg of microsomal proteins were then treated with 30 units of calf intestinal alkaline phosphatase (TaKaRa) at 37°C for 2 h. The reaction was stopped adding 3× SDS gel loading buffer.

For immunoblotting of SDS-PAGE gels without Phos-tag, total proteins were extracted, separated on 6, 7.5, or 10% SDS-PAGE gels, and blotted onto PVDF membranes, as described previously (Inada et al., 2004). Anti-RPT2, anti-PHOT1, and horseradish peroxidase (HRP)-conjugated anti-rabbit IgG antibodies were prepared as described previously (Haga et al., 2015). Anti-PHOT2 antiserum was produced in rabbit using 10× His-tagged PHOT2 products incorporating residues 294-474 as the antigen. HRP activity was detected with the super-signal west femto maximum sensitivity substrate (Thermo Scientific) and the Image Quant LAS4000 Mini device (GE Healthcare). As a loading control, the protein-blotted membranes were stained using the Pierce reversible protein staining kit (Thermo Scientific). The results were confirmed using independent samples. For the statistical analysis, the signal intensities of the protein bands were quantified with Fiji software (Schindelin et al., 2012).

### In vitro Phosphorylation Assay

In vitro phosphorylation assays of a 120-kDa protein in microsomal membranes were performed as described previously (Sakai et al., 2000), with some modifications. Briefly, microsomal membrane pellets were obtained from approximately 650 two-day-old etiolated transgenic seedlings of *pMDC7-PRT2* and *pER8,* which were grown on the half-strength OS agar medium with or without 10 µM estradiol. The pellets were resuspended in 30 μL of phosphorylation buffer (50 mM Tris-MES, pH 7.5, 5 mM MgSO4, 150 mM NaCl, 1 mM EGTA, 1 mM DTT, 0.5% Triton X-100, and a protease inhibitor mixture [Complete Mini EDTA-free; Roche Diagnostics]) by pipetting. All manipulations were performed at 4°C under a dim red safelight.

Twenty micrograms of microsomal extract were diluted to a final volume of 9 μL in phosphorylation buffer and γ-^32^P-ATP was added to a final concentration of 200 µM (specific activity, 2.5 Ci/mmol). The membrane extracts were then incubated for 2 min at 30°C and irradiated with blue light at 10 μmol m^−2^ s^−1^ for 16 min. Dark control samples were mock-irradiated. After the irradiations, the samples were mixed with an equal volume of 2× SDS gel loading buffer to stop the reaction. Ten micrograms of each sample were then electrophoresed on an a 6% SDS–PAGE gel. Gels were dried and then autoradiographed by exposure to X-ray film. The images of these films were recorded with a scanner (ES8500; Epson) and the signal intensities of the 120 kDa protein were quantified with Fiji software (Schindelin et al., 2012).

### Treatments of Protein Phosphatase Inhibitors

Cantharidin and okadaic acid (CN and OKA: Fujifilm Wako) were prepared as 50 mM and 1 mM stock solutions, respectively, in dimethyl sulfoxide (DMSO). The treatments with these chemicals were performed as described previously (Sullivan et al., 2019) with brief modifications. Briefly, the aerial portions of the hypocotyls were prepared from two-day-old etiolated seedlings with a blade under safe green light conditions. Red light-irradiation at 10 µmol m^−2^ s^−1^ for 2 min was performed prior to preparation of the segment. The segments were then dipped in half strength OS solution containing 30 µM CN, 1 µM OKA or an equivalent volume of DMSO and vacuum infiltrated for 15 min. After a subsequent incubation under darkness for 105 min at 60 rpm, the segments were irradiated with blue light at 0.1 µmol m^−2^ s^−1^ for 30 min, immediately harvested with forceps, and frozen with liquid nitrogen.

### GUS Histochemical Analysis

Transgenic seedlings carrying the *RPT2pro:GUS* gene were stained with 5-bromo-4-chloro-3-indolyl-β-glucuronide (X-Gluc), as described previously (Nagashima et al., 2008). Seedling images were obtained with an MZ-16FA /DFC500 digital stereomicroscope (Leica, http://www.leica-microsystems.com).

### VENUS Imaging

VENUS fluorescence was visualized with an MZ-16FA/DFC500 digital stereomicroscope with the YFP filter (Leica). Confocal fluorescence images were recorded as described previously (Haga et al., 2015). The VENUS signal intensity was quantified with Fiji software (Schindelin et al., 2012). To prepare hypocotyl cross sections, the *rpt2* mutant transformed with a *RPT2pro:RPT2-VENUS* gene were irradiated with blue light at 10 µmol m^−2^ s^−1^ for 6 h, subsequently mounted in 2% agarose and hand-sectioned with a blade.

### Transcriptional Analysis by qRT-PCR

Total RNA was extracted using a RNeasy kit (QIAGEN). Quantitative RT-PCR was carried out using a PCR system (StepOne; Applied Biosystems) and the Luna One-Step RT-qPCR Kit (NEB) in accordance with the manufacturer’s protocol. Triplicate PCR reactions were performed in each case and three biological independent samples were used for each gene. The primers used are listed in Supplemental Table S2 and *18S rRNA* was amplified as an internal standard.

### Measurement of Phototropic Curvature

Phototropic curvatures of hypocotyls were measured on agar medium in 0.2 ml tubes using the “advanced method” described previously (Haga et al. 2012; Haga and Kimura, 2019).

## Accession Numbers

The sequence data for this article can be found in the Arabidopsis Genome Initiative or the EMBL/GenBank data libraries under the following accession numbers: *RPT2* (AT2G30520), *PHOT1* (AT3G45780), *NPH3* (AT5G64330), *PHOT2* (AT5G58140) and *UBQ3* (AT5G03240).

## Supplemental Data

**Supplemental Figure 1.** Time course analysis of continuous light-induced phototropism in the *phot1 phot2* double mutants transformed with a *35Spro:PHOT1^LOV1Cys39Ala^* gene.

**Supplemental Figure 2.** Time course analysis of phot1 autophosphorylation in tube-grown seedlings.

**Supplemental Figure 3.** The effect of phosphatase on the *rpt2*-induced mobility shift of PHOT1 during Phos-tag SDS-PAGE.

**Supplemental Figure 4.** Evaluation of anti-PHOT2 antibody.

**Supplemental Figure 5.** Binding activity of phot2 to RPT2.

**Supplemental Figure 6.** Characterization of the *rpt2* mutants transformed with a *pMDC7-RPT2* construct.

**Supplemental Figure 7.** Suppression of phot1 dephosphorylation by protein phosphatase inhibitors.

**Supplemental Figure 8.** Effect of the proteasome inhibitor MG132 on RPT2 protein expression in wild-type seedlings under red light conditions.

**Supplemental Figure 9.** Phenotypes of the *35Spro:UBQ3* transgenic lines.

**Supplemental Table 1.** Gene-specific primers used for construction.

**Supplemental Table 2.** Gene-specific primers used for qRT-PCR.

## Acknowledgments

We thank the Arabidopsis Biological Resource Center for providing the *phot1* (Salk146058) and *phot2* (Salk142275) seeds. We also thank Professor John Christie (University of Glasgow, Glasgow, UK) for kindly providing the seeds of the *35Spro:PHOT1^LOV1C39A^* transgenic line, Professor Nam-Hai Chua (Rockefeller University, New York, NY) for generously donating the pMDC7 vector, and Dr. Masatoshi Yamaguchi (Saitama University, Saitama, Japan) and Professor Taku Demura (Nara Institute of Science and Technology) for generously supplying the pH35GS binary vectors. This work was supported by the Japan Society for the Promotion of Science (KAKENHI 22570058, 25120710, 16H01231, 17H03694, 18K19329 to T.S.).

## Author Contributions

T.K, T.T.-M. and T.S. designed and conducted most of the research, and analyzed the data. H.I., K.O. and K.I. designed and performed the phot1 in vitro phosphorylation assay. T.K. and T.S. wrote the article.

